# DAGLα/β, 2-AG release, and Parkinson’s Disease: Exploring a causal link

**DOI:** 10.1101/2025.04.17.649373

**Authors:** Patrick Doherty, Gareth Williams

## Abstract

The diacylglycerol lipases, DAGLα and DAGLβ, hydrolyse diacylglycerol (DAG) to produce 2-arachidonoylglycerol (2-AG), a key endocannabinoid (eCB) and CB_1_/CB_2_ receptor ligand. While DAGLα is well established as a regulator of CB_1_-dependent synaptic plasticity, recent studies have identified *DAGLB* mutations as a cause of autosomal recessive early-onset Parkinson’s disease (PD). Here, we present a comprehensive analysis of DAGLβ mRNA expression, demonstrating its co-expression with DAGLα mRNA predominantly in excitatory neurons throughout the adult nervous system. We see no evidence for enrichment of the DAGLs or CB_1_ transcripts in the striatum or in dopaminergic neurons. We discuss these findings within a review of recent literature that points to a wider involvement of the eCB system in PD. Notably, DAGLα-dependent 2-AG release at synapses relies on α-synuclein function—a protein central to PD pathophysiology—implicating both DAGLs in PD and pointing to widespread disruption in 2-AG release. Consistent with this, substantial reductions in 2-AG levels have been reported in the cerebrospinal fluid (CSF) of PD patients. Depression, a major non-motor symptom of PD, often precedes the onset of motor deficits by several years. Human and mouse genetic studies suggest that reduced DAGL activity may contribute to depression by impairing 2-AG-mediated CB_1_ receptor signalling, which is crucial for synaptic plasticity, stress resilience, and mood regulation. These findings point to a potential causal link between DAGL dysfunction and the non-motor symptoms in PD.

## Introduction

The diacylglycerol lipases, DAGLα and DAGLβ, are enzymes that hydrolyse diacylglycerol (DAG) in the lipid bilayer to produce 2-arachidonoylglycerol (2-AG), the primary endocannabinoid (eCB) ligand for CB_1_ and CB_2_ cannabinoid receptors [1]. They also have functions beyond cannabinoid signalling; in this context, hydrolysis of 2-AG by monoacylglycerol lipase (MAGL) generates arachidonic acid (AA), which is involved in the production of pro-inflammatory eicosanoids (e.g., prostaglandins, leukotrienes), implicating DAGLs in the regulation of inflammatory lipid signalling [2]. DAGLα knockout mice show an approximate 80% reduction in 2-AG levels across multiple brain regions, including the whole brain, amygdala, cerebellum, hippocampus, striatum, prefrontal cortex, and hypothalamus [3-7]. In contrast, DAGLβ knockout mice show a 50% reduction in whole-brain 2-AG, but no significant changes in the cortex, cerebellum, hippocampus, or striatum, with only a small but significant reduction in the hypothalamus [3-5]. Analysis of individual cell types cultured from knockout mice show DAGLα to be solely responsible for 2-AG steady-state levels in neurons, largely responsible for this in astrocytes, with no role in microglia. In contrast, DAGLβ was the dominant enzyme in microglia, played a minor role in astrocytes, and had no clear function in neurons [8].

DAGLα and DAGLβ share a conserved amino-terminal four-transmembrane domain and an intracellular catalytic domain, but DAGLα has a significantly longer carboxy-terminal tail [2]. In the adult, DAGLα appears to be the “synaptic” enzyme, localised to dendritic spines in the hippocampus, cortex, and cerebellum [9, 10]. This likely depends on interactions between its unique tail and Homer proteins [11], which can regulate eCB mediated plasticity [12] and are essential for organising receptor complexes at postsynaptic densities [13]. At the membrane, 2-AG synthesis is initiated by depolarisation and activation of phospholipase C (PLC) via both ionotropic receptors (e.g., NMDA receptors) and metabotropic receptors (e.g., mGluRs) [14]. Two distinct mechanisms of 2-AG production have been described: one that is strictly calcium-dependent, and another that is initiated by G-protein coupled receptors and facilitated by calcium [14]. These findings suggest the existence of functionally distinct pools of DAGLα. In this context, cycling of DAGLα between the plasma membrane and an intracellular endosomal compartment generates separate pools of the enzyme and dynamically regulates its availability at the postsynaptic membrane [15]. After synthesis, evidence suggests that 2-AG is released from cells associated with microvesicles [16], with the vesicles most likely shed from the membrane by a budding mechanism [17].

Once released, 2-AG activates presynaptic CB_1_ receptors on excitatory and/or inhibitory synapses modulating both short- and long-term synaptic plasticity, including depolarisation-induced suppression of excitation (DSE) and inhibition (DSI) [18]. This form of plasticity is abolished throughout the brain when DAGLα is knocked out but is unaffected by DAGLβ knockout [3-5]. At synapses, the activity of 2-AG is tightly regulated by enzymatic hydrolysis by MAGL [19]. Additionally, α/β-hydrolase domain-containing 6 (ABHD6), an intracellular enzyme found in the postsynaptic compartment, and ABHD12, an extracellular enzyme associated with microglia, can contribute to 2-AG hydrolysis [8].

DAGLβ is enriched in macrophages and dendritic cells, and as mentioned is the major source of 2-AG in microglia, the resident immune cells in the brain. In these cells, DAGLβ works hand in glove with MAGL to synthesise AA that serves as a precursor for pro-inflammatory eicosanoids like prostaglandins, crucial for initiating and sustaining often detrimental inflammatory responses [20]. Microglia also express CB_2_ receptors, and activation of these exerts anti-inflammatory effects, reducing pro-inflammatory cytokine release and promoting microglial motility and phagocytosis [21]. This can be neuroprotective in conditions like neurodegeneration and neuroinflammation.

Parkinson’s disease (PD), the second most common neurodegenerative disorder after Alzheimer’s, affects 1–2% of people over 65. Its primary pathological feature is the loss of dopaminergic neurons (DANs) in the substantia nigra leading to dopamine depletion in the nigrostriatal pathway and characteristic motor symptoms with a key feature being the accumulation of α-synuclein aggregates in Lewy bodies in nigral DANs and other neurons [22, 23]. Estimates suggest that 5-10% of PD cases are due to monogenic mutations in genes like *SNCA* (α-synuclein), *LRRK2, VPS35, PRKN* (parkin), *PINK1*, and *DJ-1* and these mutations often lead to early-onset forms of the disease [24].

A study by Liu et al., recently identified biallelic mutations in *DAGLB* as a cause of autosomal recessive early-onset PD [25]. They reported four homozygous mutations in six individuals from four families, all of whom developed symptoms before 40 and initially responded well to levodopa, with PET confirming impaired dopamine transmission in the striatum. Four distinct mutations were documented, (two missense and two within introns) with all four reported to affect the formation or stability of the DAGLβ protein arguing for a loss of function phenotype. Later research has extended these findings beyond the Chinese population. Tesson et al., identified a homozygous missense variant in a consanguineous North African patient with early-onset PD [26]. This variant affects a conserved amino acid in the protein’s catalytic domain, near one of the mutations reported by Liu et al. (the missense mutations are shown schematically in Fig. 1). Notably, *DAGLB* mutations are extremely rare, and the same study noted no pathogenic variants in sporadic late-onset PD cases of European descent.

**Figure 1.**
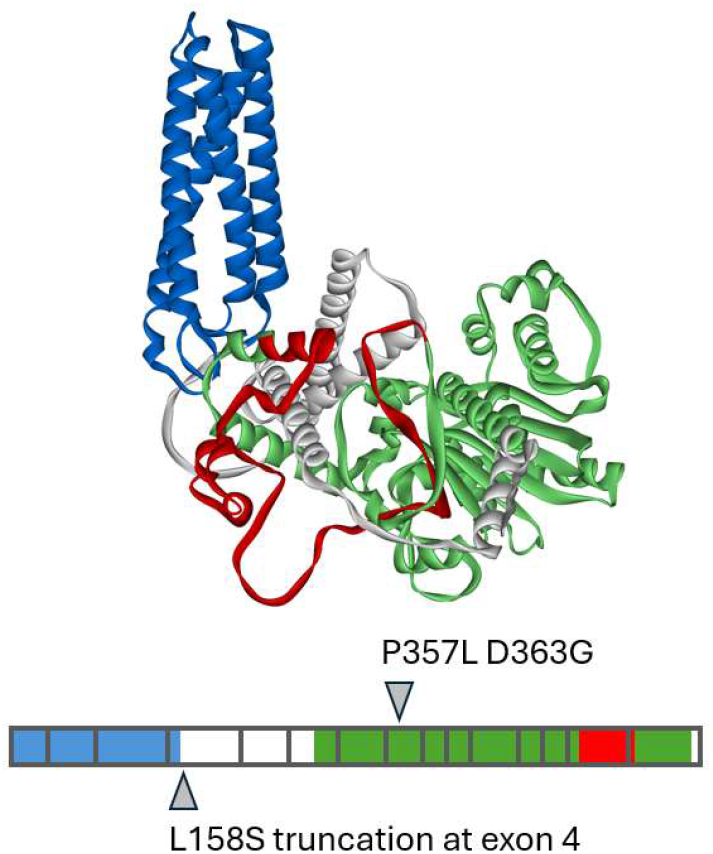
DAGLb mutations associated with PD susceptibility. At top the AlphaFold structure for DAGLb in humans is shown. The four-helix transmembrane domain is coloured blue, the hydrolase catalytic domain is in green and the regulatory loop in red. The exon structure is shown below with the positions of the three coding variants.

In this study, we provide a comprehensive analysis of DAGLβ mRNA expression, demonstrating its co-expression with DAGLα mRNA primarily in excitatory neurons throughout the adult mouse and human brain. We find no evidence of DAGLβ mRNA enrichment in the striatum or in nigral DANs. We discuss these findings within a wide review of the recent literature and discuss the possibility that loss of DAGLβ or DAGLα function may disrupt widespread neural network activity and contribute to depression in PD.

## Methods

### Single cell RNAseq data

Single cell RNAseq FKPM data for human DAMs and motor neurons was obtained from NCBI GEO [27] under the series accession GSE76514 [28]. FPKM counts corresponding to the given genes were compared between the 27 dopaminergic neurons and 24 motor neurons cells. One sample was dropped as it was 7.5 standard deviations above the mean. Mouse single cell RNAseq FKPM data for Th-eGFP+ sorted neurons was obtained from NCBI GEO under the accession GSE108020 [29]. The data comprises cell counts for 473 cells with varying TH counts. The cells were categorised as DANs if their TH counts exceed 100. The averages in the two classes are then 1396.57+/-141.58 and 9.59+/-1.49

### Allan Brain Atlas anatomic structure data

RNAseq data for two human donors across 121 anatomically distinct brain regions was downloaded from the Allen Brain portal human.brain-map.org/static/download. Cell count data was combined from the two donors and averaged over striatal and non-striatal regions.

### Allan Brain Atlas cell type data

Human multiple cortical areas and Mouse multiple cortical areas and the hippocampal formation SMART-seq [30] single cell data were downloaded from the Allen Brain Atlas cell type portal portal.brain-map.org/atlases-and-data/rnaseq. The 2D tSNE (for the Human data) and UMAP (for the Mouse data) coordinates for the cells were downloaded from the same portal and merged with the cell count data for the genes under consideration in the R scripting environment. The relative expression levels across the main cell types were calculated by averaging the trimmed mean expression data. The Human trimmed mean data consists of mean counts for 120 clusters of the 49,495 samples. These clusters were in turn grouped into 9 groups corresponding to: excitatory neurons, inhibitory neurons, astrocytes, endothelial cells, vascular and leptomeningeal cells, oligodendrocytes, pericytes, oligodendrocyte precursors, and microglia. The Mouse data comprises 382 clusters and these were grouped into 8 groups corresponding to: excitatory neurons, inhibitory neurons, astrocytes, endothelial cells, vascular and leptomeningeal cells, oligodendrocytes, smooth muscle cells, and microglia.

## Results and Discussion

*DAGLβ, an underappreciated role in neuronal function*. An obvious question arising from the identification of *DAGLB* mutation as a cause of PD is whether it is expressed in nigral DANs, given their vulnerability in PD. Using single-cell RNAseq, DAGLβ has not only been found in both human and mouse nigral DANs, but surprisingly it was 10-fold more abundant than DAGLα in the human cells [25]. In this study, we investigate whether DAGLβ expression in nigral DANs is unusual compared to other neuronal types. Using the same GSE76514 dataset obtained from human neurons [28], we confirmed that DAGLβ transcripts are approximately ten times more abundant than DAGLα transcripts in human nigral DANs; however, large standard errors associated with both means indicate substantial variability, making the precise magnitude of this difference uncertain (not shown). More pertinently, in the same dataset overall DAGLβ transcript levels do not differ significantly between nigral DANs and motor neurons. With the relative levels in the human neurons being 3.24+/-0.90 (DAN) versus 3.39+/-0.57 (motor neurons) (here Grubbs’ test was applied to the data and one clear outlier dropped from the DAN set). This result demonstrates that nigral DANs do not uniquely express high levels of this transcript.

To gain further insights, we analysed RNAseq data from the Allen Brain Atlas for human and mouse brain cells. A striking initial observation is that DAGLβ mRNA is clearly co-expressed with DAGLα mRNA in human and mouse cortical neurons, largely but not exclusively within the clusters of excitatory neurons (Fig. 2). In particular, the overall Pearson correlation in expression between the two transcripts is 0.62 (p < 3.1e-14) in the human cells and 0.59 (p < 7.9e-38) in mouse cells. However, note also the relatively exclusive expression of DAGLβ in microglia and DAGLα in astrocytes that is obvious in the human cell clusters. In terms of average levels of mRNA transcript expression, in human striatal versus non-striatal tissue (see Fig. 3a) there is a 2-3-fold enrichment of DAGLα and DAGLβ in non-striatal tissue compared to the striatum. In both regions, DAGLα is expressed at higher levels than DAGLβ, with CB_1_ equally expressed. A separate study used single-cell RNAseq to identify distinct dopaminergic neuronal subpopulations in developing mouse brains [29]. We interrogated their data (GSE108020, [29]) defining TH+ cells based on a transcript count threshold, we find similar DAGLα and DAGLβ levels in both TH+ and TH− populations. However, CB_1_ expression is lower in the TH+ population (Fig. 3b).

**Figure 2.**
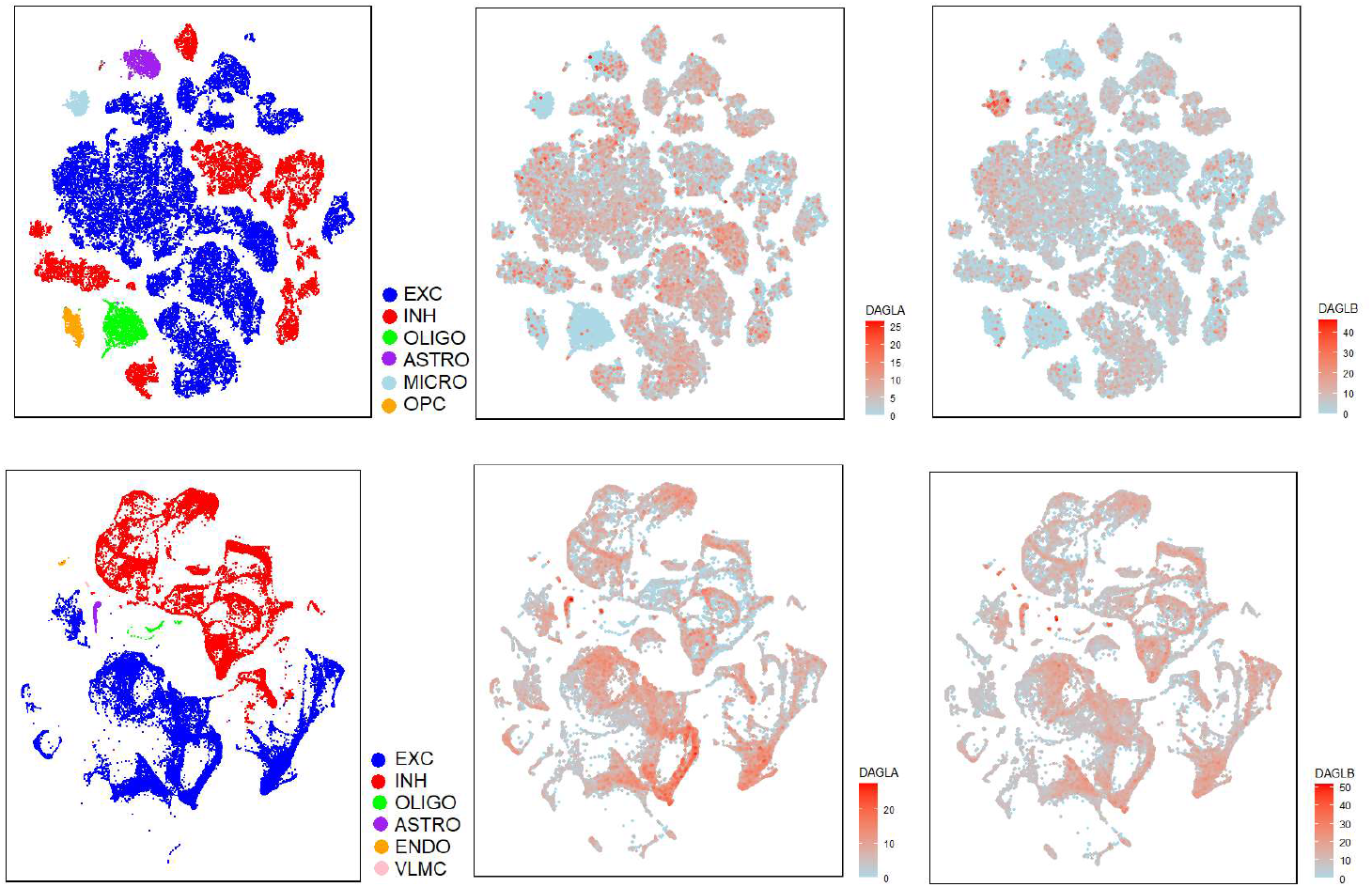
DAGL distribution across brain cell types in human and mouse. The top panel shows the tSNE projection of 49,495 cells from multiple human cortical areas. The cells cluster into distinct classes based on SMARTseq gene expression and these are indicated at the left. It is clear that both DAGLα and DAGLβ show wide expression, mostly in the excitatory cells. Interestingly DAGLb shows a market expression in the microglial cluster (light blue). The bottom panel shows the UMAP projection of 77,000 cells from cortical and hippocampal areas of the mouse brain. The same picture emerges as with the human data.

**Figure 3.**
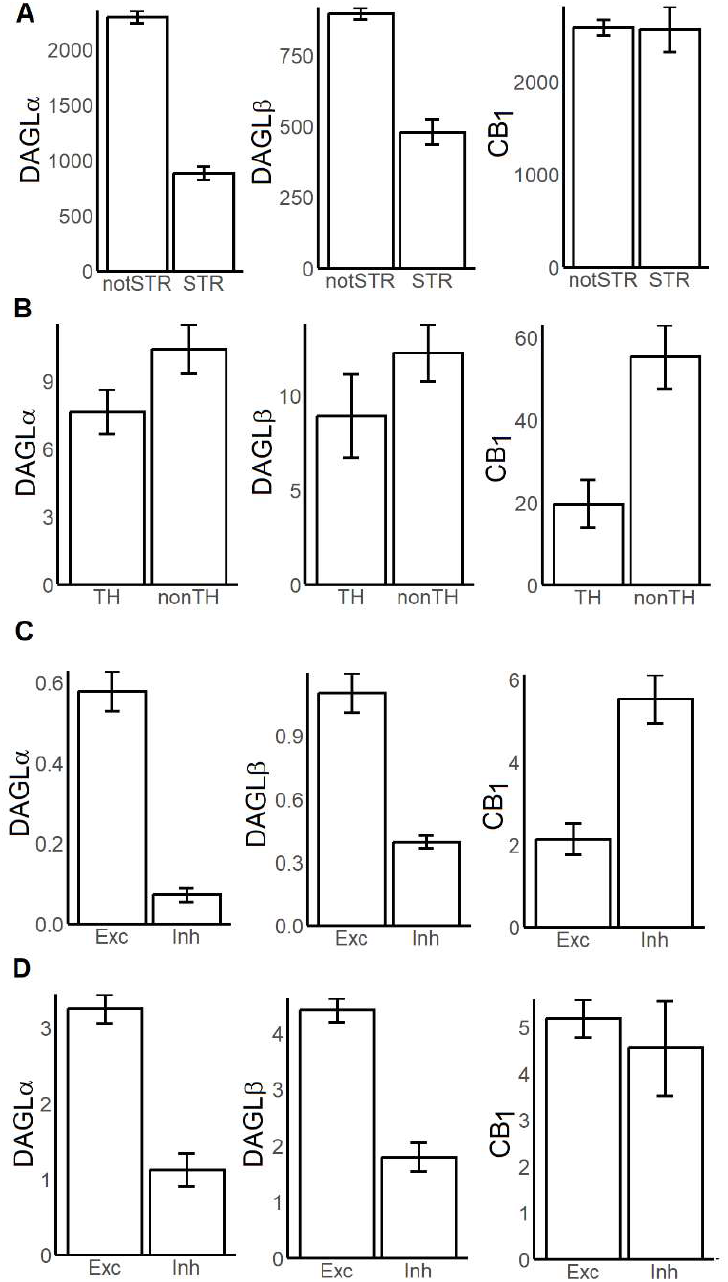
Relative expression of DAGLα, DAGLβ and CB_1_ in human and mouse neurons. In A the expression levels are compared between striatum and all other brain regions from RNAseq expression data from human brains from the Allen Brain Atlas. The striatum is characterised by a relatively high expression of the TH dopaminergic neuron marker (relative levels of TH 176.29+/-25.90 and 20.77+/-0.87 p < 1.8e-05). Data from single cell RNAseq expression of 473 mouse brain cells sorted based on Th-eGFP+ is shown in B. Here the TH and nonTH cells were defined based on a threshold count of TH expression (a threshold of 100 results in TH levels of 1396.57+/-141.58 v 9.59+/-1.49 p < e-16 in the two categories). As in A there is a moderate drop in both DAGL isoforms in the TH expressing cells. Interestingly CB_1_ appears to be higher in nonTH cells. C and D show the Human and Mouse single cell cortical neuron RNAseq data expression levels. Data is based on cluster means in 121 (HUMAN) and 382 (MOUSE) clusters of the full data set. These clusters in turn segregate into inhibitory and excitatory super clusters for which the data is given. Consistently, the DAGL enzymes are higher in excitatory neurons with hardly any expression in the Human inhibitory cells.

Finally, we examined DAGLα, DAGLβ, and CB_1_ mRNA expression in the excitatory and inhibitory neuronal clusters human and mouse cortex illustrated in Fig. 2. In humans, DAGLα and DAGLβ are expressed at similar levels in clusters of excitatory neurons, but at lower levels in clusters of inhibitory neurons. In both cases DAGLβ transcripts were more abundant than DAGLα transcripts, but only by 2-3-fold. The CB_1_ transcript is readily detected in both excitatory and inhibitory neuronal clusters, with higher levels in the latter (Fig. 3c). A similar pattern is observed in the mouse cortex, with some minor differences. For example, DAGLα and DAGLβ are again expressed at higher levels in excitatory versus inhibitory neuronal clusters, but in this case at similar levels to each other (Fig. 3d). Likewise, CB_1_ transcript levels were similar between the two neuronal populations.

It is important to note that protein abundance is influenced not only by transcript levels but also by translational efficiency and protein turnover dynamics, which may differ between DAGLα and DAGLβ. Nonetheless, at first glance, the observation of widespread presence of DAGLβ transcripts in neurons appears at odds with the marginal effects of DAGLβ knockout on 2-AG levels reported in most brain regions and neuronal cultures from knockout mice. However, these studies measure steady-state 2-AG levels, not the signalling pools of 2-AG and notably MAGL inhibition in DAGLα knockout mice can lead to a substantial DAGL-dependent increase in brain 2-AG levels, suggesting that DAGLβ may play a more substantial role in 2-AG synthesis that is masked by rapid hydrolysis [6, 8]. In line with this, the task learning associated DAGLβ-dependent 2-AG release from nigral DANs was only seen in the dorsal striatum following MAGL inhibition [25]. Furthermore, both enzymes have been shown to cooperate in generating signalling pools of 2-AG in neural stem cells [31] and cultured hippocampal neurons [32].

In summary, DAGLα and DAGLβ appear to be co-expressed in neurons, though their relative abundance and subcellular distribution likely vary. Importantly, there is nothing unique about expression of DAGLβ, or indeed DAGLα or CB_1_ in nigral DANs or the striatum to suggest increased vulnerability to loss of eCB signalling resulting in a PD motor pathology. The findings are nonetheless significant as they offer a plausible mechanism by which DAGLβ loss in PD patients might compromise widespread neural network activity. Also, given their co-expression, they also provide a plausible explanation for a potential compensatory role for DAGLα in mitigating the effects of DAGLβ loss in transgenic mice.

### DAGLβ knockout is not associated with development of PD pathophysiology in mice

The ability of mice to balance on a rotating rod is often used as a sensitive measurement of inherent motor function [33]. Their ability to stay on the rod improves with training with striatal dopaminergic modulation playing a supportive, but not exclusive, role in consolidating the learned motor skills [34]. DAGLβ knockout mice are not impaired in their inherent ability to balance on the rotarod, nor their ability to learn the task. In addition, there is no loss of nigral DANs or other PD-like features (e.g., neuroinflammation, axon swelling, or mitochondrial deformation) in DAGLβ knockout mice at up to 20 months of age [25]. Similarly, the same study reported no impairment of baseline function in mice that had DAGLβ selectively knocked down in nigral DANs at 3 months of age, nor PD like pathology in these mice at up to 12 months, ruling out a simple developmental compensation.

However, the authors demonstrated that the selective knockdown of DAGLβ in the adult nigral DANs compromised the mice’s ability to learn the rotarod task, and this was associated with a reduction in the steady state levels of 2-AG in the dorsal striatum and a blunting of a learning associated increase in 2-AG. The observation of a *normal* baseline rotarod performance and *only* a learning deficit is very unusual for PD models but could reflect an early stage of the disease. The relationship of this phenotype to PD and the absence of a similar phenotype in DAGLβ knockout mice call for further investigation.

### Synucleins are required for 2-AG release at synapses

α-Synuclein (*SNCA*) is unique among Parkinson’s-related genes, being both a causative factor in rare familial cases and central to the pathology of all forms of the disease. *SNCA* mutations, gene duplications, and common variants increase disease risk by promoting α-synuclein misfolding and aggregation into Lewy bodies [23]. While aggregation of α-synuclein in Parkinson’s disease suggests a toxic gain-of-function mechanism, the concomitant sequestration of the physiological pool into aggregates may also contribute via loss-of-function; these mechanisms are not mutually exclusive and may act in concert.

Although subtle defects in the nigrostriatal dopamine system have been reported in α-synuclein knockout mice [35], due to functional redundancy among α-, β-, and γ-synucleins, the physiological roles of this protein family are best revealed in triple knockout models, where compensatory mechanisms are eliminated. Triple knockout of α-, β-, and γ-synucleins reveals that while synucleins are not essential for neuronal development or survival, they play key roles in maintaining presynaptic function and dopamine homeostasis. Triple knockout mice exhibit enhanced dopamine release and turnover in the dorsal striatum, reduced dopamine stores, and age-dependent synaptic dysfunction without loss of neurons. Synucleins regulate excitatory synapse size and preserve long-term synaptic integrity, suggesting their disruption may contribute to the regional vulnerability and progression of Parkinson’s disease [36-38].

Remarkably, triple transgenic knockout mice lacking all synucleins show impaired eCB release leading to deficits in long-term depression (LTD) and DSI at excitatory and inhibitory synapses in the striatum and hippocampus [39]. This defect originates postsynaptically, as CB_1_ receptor function remains intact. By directly loading neurons with the eCBs, researchers found that synucleins are required for normal 2-AG release, but not its synthesis. Rescue experiments revealed that α-synuclein’s membrane-binding ability is essential for this function, with PD-associated A30P mutants unable to restore eCB release. Synucleins act as SNARE chaperones, promoting SNARE complex assembly during vesicular exocytosis and recycling and inactivation of synaptobrevin-2 blocked eCB release and impaired eCB-dependent plasticity. However, 2-AG release appears to occur via vesicle shedding rather than conventional exocytosis [17] leaving the precise mechanism affected by synaptobrevin-2 blockade unresolved. Intriguingly, the surface pool of DAGLα is dynamically regulated, undergoing endocytosis and recycling from an intracellular endosomal compartment to the postsynaptic membrane [15]. If 2-AG release is coupled to this recycling process it offers a potential mechanism that might be dependent on synuclein/SNARE function. Nevertheless, irrespective of the mechanism, the collaboration between synucleins and SNAREs in postsynaptic eCB release highlights a pathway linking DAGLα to PD, potentially through widespread dysregulation of eCB signalling.

### Reduced CSF 2-AG Levels in Parkinson’s Disease Patients

While changes in signalling lipids, including 2-AG, have been extensively studied in animal models, there is a notable lack of data on 2-AG levels in PD patient samples [40]. However, a recent study reported a highly significant 39% reduction in cerebrospinal fluid (CSF) 2-AG levels in PD patients compared to healthy controls [41]. Notably, this reduction was seen in PD patients both with and without levodopa-induced dyskinesia, suggesting that 2-AG depletion may be an early feature of PD pathophysiology.

In PD, it is has recently been estimated that 35-45% of nigral DANs are lost by the time motor symptoms become noticeable [42], such a loss alone could not account for the substantial reduction in CSF 2-AG levels. Moreover, neurodegeneration-associated inflammation would be expected, if anything, to increase 2-AG production as part of the brain’s response to injury [43]. Therefore, the observed reduction in 2-AG points to a more systemic dysfunction of eCB signalling. The underlying cause of this reduction is still unclear, highlighting the need for further investigation into how 2-AG dysregulation correlates with disease progression.

### DAGLA and DAGLB: Genetic Links to Depression with Implications for Parkinson’s Disease

Depression is a common non-motor symptom of Parkinson’s disease (PD), involving neural circuits distinct from those responsible for motor dysfunction. While motor symptoms are primarily linked to degeneration in the nigrostriatal pathway, PD-related depression is associated with alterations in the mesolimbic and mesocortical dopaminergic pathways, as well as disruptions in serotonergic and noradrenergic systems [44, 45]. Depression can be manifest several years before the development of classical motor symptoms, and indeed some evidence suggests that early disturbances in these mood-regulating circuits may contribute to the later development of motor symptoms, reinforcing the intricate interplay between emotional and motor pathways in PD [14].

The observation that 2-AG release may be compromised by α-synuclein pathology raises the possibility that this could provide a mechanistic link between DAGLα, DAGLβ, and PD. Notably, a depression- and anxiety-like phenotype has been reported in independent DAGLα knockout mouse lines [6, 7, 46]. In humans, a cross-disorder GWAS meta-analysis involving over 284,000 cases and 508,000 controls identified *DAGLA* as the top endocannabinoid gene associated with psychiatric risk. *DAGLA* showed the strongest association with mental disorder risk, including major depressive disorder, suggesting that dysfunction in this key 2-AG–synthesising enzyme may contribute to the genetic architecture of depression and related conditions [47].

When findings from an independent GWAS meta-analysis are integrated with Mendelian Randomisation analyses using protein quantitative trait loci and functional biological data, DAGLα again emerges as a top candidate associated with depression and suggested by the authors as a promising therapeutic target. Interestingly, while *DAGLB* does not reach genome-wide significance in GWAS data alone, it is also identified as significantly associated with depression when these multi-omics approaches are combined [48].

Clinically, the CB_1_ receptor antagonist rimonabant—originally developed for obesity treatment— effectively mimics reduced DAGLα activity by blocking 2-AG signalling. However, rimonabant was withdrawn from clinical use due to severe psychiatric side effects, including depression, anxiety, insomnia, and increased risk of suicidal ideation [49]. These adverse effects likely reflect disruptions in monoaminergic neurotransmission (e.g., serotonin and noradrenaline), highlighting the critical role of the endocannabinoid system in mood regulation. Together, these findings offer a compelling mechanistic framework linking DAGLα and DAGLβ dysfunction to the non-motor symptoms of PD, particularly depression and anxiety.

## Concluding remarks and future directions

Our data analysis provides the first clear evidence of widespread co-expression of DAGLβ and DAGLα in neurons across both the mouse and human brain, highlighting an under-appreciated role for DAGLβ in contributing to neuronal 2-AG synthesis. This suggests that DAGLB mutations in early-onset Parkinson’s disease (PD) may cause widespread disruption of neural networks, rather than limited dysfunction within nigral DANs. The essential role of synucleins in DAGLα-dependent 2-AG release further supports the idea of broad eCB system dysregulation in PD.

With respect to impaired 2-AG release and PD symptoms, loss of CB1 signalling can cause depression-like behaviour in mice and is associated with depression in humans. GWAS and related studies have identified both DAGLA and DAGLB as genetic risk factors. These findings strongly support a link between the DAGLs, 2-AG release, and major non-motor symptoms in PD. The intriguing observation of reduced 2-AG levels in the cerebrospinal fluid of PD patients further reinforces this connection. However, the mechanisms driving this reduction, and its relationship to disease progression, remain unclear and warrant further investigation.

Finally, by way of analogy, rare TREM2 variants are linked to early-onset Alzheimer’s disease, with dysfunctional TREM2 impairing microglial responses, leading to defective amyloid-β clearance, increased neuroinflammation, and accelerated neurodegeneration [50-52]. This serves as a potent reminder that exploring the role of DAGL dysfunction beyond neurons, particularly in microglia, could reveal previously unappreciated facets of PD pathology and suggest novel therapeutic strategies.

## References

1. Bisogno, T., et al., Cloning of the first sn1-DAG lipases points to the spatial and temporal regulation of endocannabinoid signaling in the brain. J Cell Biol, 2003. 163(3): p. 463–8.

2. Reisenberg, M., et al., The diacylglycerol lipases: structure, regulation and roles in and beyond endocannabinoid signalling. Philos Trans R Soc Lond B Biol Sci, 2012. 367(1607): p. 3264–75.

3. Gao, Y., et al., Loss of retrograde endocannabinoid signaling and reduced adult neurogenesis in diacylglycerol lipase knock-out mice. J Neurosci, 2010. 30(6): p. 2017–24.

4. Tanimura, A., et al., The endocannabinoid 2-arachidonoylglycerol produced by diacylglycerol lipase alpha mediates retrograde suppression of synaptic transmission. Neuron, 2010. 65(3): p. 320–7.

5. Yoshino, H., et al., Postsynaptic diacylglycerol lipase mediates retrograde endocannabinoid suppression of inhibition in mouse prefrontal cortex. J Physiol, 2011. 589(Pt 20): p. 4857–84.

6. Shonesy, B.C., et al., Genetic disruption of 2-arachidonoylglycerol synthesis reveals a key role for endocannabinoid signaling in anxiety modulation. Cell Rep, 2014. 9(5): p. 1644–1653.

7. Jenniches, I., et al., Anxiety, Stress, and Fear Response in Mice With Reduced Endocannabinoid Levels. Biol Psychiatry, 2016. 79(10): p. 858–868.

8. Viader, A., et al., A chemical proteomic atlas of brain serine hydrolases identifies cell type-specific pathways regulating neuroinflammation. Elife, 2016. 5: p. e12345.

9. Katona, I., et al., Molecular composition of the endocannabinoid system at glutamatergic synapses. J Neurosci, 2006. 26(21): p. 5628–37.

10. Yoshida, T., et al., Localization of diacylglycerol lipase-alpha around postsynaptic spine suggests close proximity between production site of an endocannabinoid, 2-arachidonoyl-glycerol, and presynaptic cannabinoid CB1 receptor. J Neurosci, 2006. 26(18): p. 4740–51.

11. Jung, K.M., et al., A key role for diacylglycerol lipase-alpha in metabotropic glutamate receptor-dependent endocannabinoid mobilization. Mol Pharmacol, 2007. 72(3): p. 612–21.

12. Roloff, A.M., et al., Homer 1a gates the induction mechanism for endocannabinoid-mediated synaptic plasticity. J Neurosci, 2010. 30(8): p. 3072–81.

13. Shiraishi-Yamaguchi, Y. and T. Furuichi, The Homer family proteins. Genome Biol, 2007. 8(2): p. 206.

14. Goldman, J.G. and R. Postuma, Premotor and nonmotor features of Parkinson’s disease. Curr Opin Neurol, 2014. 27(4): p. 434–41.

15. Zhou, Y., et al., Regulated endosomal trafficking of Diacylglycerol lipase alpha (DAGLalpha) generates distinct cellular pools; implications for endocannabinoid signaling. Mol Cell Neurosci, 2016. 76: p. 76–86.

16. Lombardi, M., et al., Extracellular vesicles released by microglia and macrophages carry endocannabinoids which foster oligodendrocyte differentiation. Front Immunol, 2024. 15: p. 1331210.

17. Straub, V.M., et al., The endocannabinoid 2-arachidonoylglycerol is released and transported on demand via extracellular microvesicles. Proc Natl Acad Sci U S A, 2025. 122(8): p. e2421717122.

18. Oudin, M.J., C. Hobbs, and P. Doherty, DAGL-dependent endocannabinoid signalling: roles in axonal pathfinding, synaptic plasticity and adult neurogenesis. Eur J Neurosci, 2011. 34(10): p. 1634–46.

19. Hashimotodani, Y., T. Ohno-Shosaku, and M. Kano, Presynaptic monoacylglycerol lipase activity determines basal endocannabinoid tone and terminates retrograde endocannabinoid signaling in the hippocampus. J Neurosci, 2007. 27(5): p. 1211–9.

20. Hsu, K.L., et al., DAGLbeta inhibition perturbs a lipid network involved in macrophage inflammatory responses. Nat Chem Biol, 2012. 8(12): p. 999–1007.

21. Komorowska-Muller, J.A. and A.C. Schmole, CB2 Receptor in Microglia: The Guardian of Self-Control. Int J Mol Sci, 2020. 22(1).

22. Surmeier, D.J., J.A. Obeso, and G.M. Halliday, Selective neuronal vulnerability in Parkinson disease. Nat Rev Neurosci, 2017. 18(2): p. 101–113.

23. Moore, D.J., et al., Molecular pathophysiology of Parkinson’s disease. Annu Rev Neurosci, 2005. 28: p. 57–87.

24. Blauwendraat, C., M.A. Nalls, and A.B. Singleton, The genetic architecture of Parkinson’s disease. Lancet Neurol, 2020. 19(2): p. 170–178.

25. Liu, Z., et al., Deficiency in endocannabinoid synthase DAGLB contributes to early onset Parkinsonism and murine nigral dopaminergic neuron dysfunction. Nat Commun, 2022. 13(1): p. 3490.

26. Tesson, C., et al., Identification of a DAGLB Mutation in a Non-Chinese Patient with Parkinson’s Disease. Mov Disord, 2023. 38(9): p. 1756–1757.

27. Edgar, R., M. Domrachev, and A.E. Lash, Gene Expression Omnibus: NCBI gene expression and hybridization array data repository. Nucleic Acids Res, 2002. 30(1): p. 207–10.

28. Nichterwitz, S., et al., Laser capture microscopy coupled with Smart-seq2 for precise spatial transcriptomic profiling. Nat Commun, 2016. 7: p. 12139.

29. Hook, P.W., et al., Single-Cell RNA-Seq of Mouse Dopaminergic Neurons Informs Candidate Gene Selection for Sporadic Parkinson Disease. Am J Hum Genet, 2018. 102(3): p. 427–446.

30. Hodge, R.D., et al., Conserved cell types with divergent features in human versus mouse cortex. Nature, 2019. 573(7772): p. 61–68.

31. Goncalves, M.B., et al., A diacylglycerol lipase-CB2 cannabinoid pathway regulates adult subventricular zone neurogenesis in an age-dependent manner. Mol Cell Neurosci, 2008. 38(4): p. 526–36.

32. Jain, T., et al., Diacylglycerol lipasealpha (DAGLalpha) and DAGLbeta cooperatively regulate the production of 2-arachidonoyl glycerol in autaptic hippocampal neurons. Mol Pharmacol, 2013. 84(2): p. 296–302.

33. Brooks, S.P. and S.B. Dunnett, Tests to assess motor phenotype in mice: a user’s guide. Nat Rev Neurosci, 2009. 10(7): p. 519–29.

34. Shiotsuki, H., et al., A rotarod test for evaluation of motor skill learning. J Neurosci Methods, 2010. 189(2): p. 180–5.

35. Abeliovich, A., et al., Mice lacking alpha-synuclein display functional deficits in the nigrostriatal dopamine system. Neuron, 2000. 25(1): p. 239–52.

36. Anwar, S., et al., Functional alterations to the nigrostriatal system in mice lacking all three members of the synuclein family. J Neurosci, 2011. 31(20): p. 7264–74.

37. Greten-Harrison, B., et al., alphabetagamma-Synuclein triple knockout mice reveal age-dependent neuronal dysfunction. Proc Natl Acad Sci U S A, 2010. 107(45): p. 19573–8.

38. Burre, J., et al., Alpha-synuclein promotes SNARE-complex assembly in vivo and in vitro. Science, 2010. 329(5999): p. 1663–7.

39. Albarran, E., et al., Postsynaptic synucleins mediate endocannabinoid signaling. Nat Neurosci, 2023. 26(6): p. 997–1007.

40. Chiurchiu, V., et al., Lipidomics of Bioactive Lipids in Alzheimer’s and Parkinson’s Diseases: Where Are We? Int J Mol Sci, 2022. 23(11).

41. Marchioni, C., et al., Endocannabinoid levels in patients with Parkinson’s disease with and without levodopa-induced dyskinesias. J Neural Transm (Vienna), 2020. 127(10): p. 1359–1367.

42. Heng, N., et al., Striatal Dopamine Loss in Early Parkinson’s Disease: Systematic Review and Novel Analysis of Dopamine Transporter Imaging. Mov Disord Clin Pract, 2023. 10(4): p. 539–546.

43. Witting, A., et al., P2X7 receptors control 2-arachidonoylglycerol production by microglial cells. Proc Natl Acad Sci U S A, 2004. 101(9): p. 3214–9.

44. Schapira, A.H.V., K.R. Chaudhuri, and P. Jenner, Non-motor features of Parkinson disease. Nat Rev Neurosci, 2017. 18(8): p. 509.

45. Chaudhuri, K.R. and A.H. Schapira, Non-motor symptoms of Parkinson’s disease: dopaminergic pathophysiology and treatment. Lancet Neurol, 2009. 8(5): p. 464–74.

46. Schuele, L.L., et al., Diacylglycerol lipase alpha in astrocytes is involved in maternal care and affective behaviors. Glia, 2021. 69(2): p. 377–391.

47. Kim, H.K., et al., Cross-disorder GWAS meta-analysis of endocannabinoid DNA variations in major depressive disorder, bipolar disorder, attention deficit hyperactivity disorder, autism spectrum disorder, and schizophrenia. Psychiatry Res, 2023. 330: p. 115563.

48. Li, Y., et al., Cross-ancestry genome-wide association study and systems-level integrative analyses implicate new risk genes and therapeutic targets for depression. Nat Hum Behav, 2025.

49. Christensen, R., et al., Efficacy and safety of the weight-loss drug rimonabant: a meta-analysis of randomised trials. Lancet, 2007. 370(9600): p. 1706–13.

50. Wang, Y., et al., TREM2 lipid sensing sustains the microglial response in an Alzheimer’s disease model. Cell, 2015. 160(6): p. 1061–71.

51. Guerreiro, R., et al., TREM2 variants in Alzheimer’s disease. N Engl J Med, 2013. 368(2): p. 117–27.

52. Qin, Q., et al., TREM2, microglia, and Alzheimer’s disease. Mech Ageing Dev, 2021. 195: p. 111438.

